# Fungus-infected *Meiogyne* flowers offer a brood site for beetle pollinators in a tripartite nursery pollination system

**DOI:** 10.1101/2024.09.19.613827

**Authors:** Ming-Fai Liu, Junhao Chen, Bine Xue, Rui-Jiang Wang, Richard M. K. Saunders

## Abstract

Fungi are widely known for their pathological impact on flowers, but some play a beneficial role in pollination. We report a case of tripartite pollination system in the flowering plant *Meiogyne hainanensis* (Annonaceae) in Hainan, China. The flowers emit a fruity scent composed of a mixture of mostly sesquiterpenes and aliphatic esters, attracting the primary beetle pollinators *Paraphloeostiba* sp. (Staphylinidae) and *Mimemodes* sp. (Monotomidae). The pollinators utilise the floral chamber as a mating ground and oviposit onto the adaxial corrugations of the inner petals. After the end of anthesis, extensive growth of filamentous fungi was observed to be restricted on these corrugated tissues. Upon hatching, the *Paraphloeostiba* and *Mimemodes* larvae consumed the fungal mycelia. ITS2 metabarcoding analysis reveals that the diet of the larvae consists of similar fungal taxa as those found on the inner petals. Both were primarily composed of ascomycete fungi such as *Fusarium*, *Penicillium* and *Cladosporium* species. The flower has an unusually long post-anthetic phase that lasts at least 21 days and up to 2 months, during which the fungus-infested petals remain arboreal, offering suitable microclimate and shelter for the broods. This is the second reported angiosperm genus that exhibits tripartite brood-site pollination in which filamentous fungi are an essential mutualistic partner.

## Introduction

Around 90% of flowering plants are pollinated by animal agents (Ollerton et al., 2011). Flower-insect interaction was hypothesised to have driven the diversification of pollinating insect lineages (Peris & Condamine, 2024). In these systems, flowers have often evolved to provide pollinators with various floral rewards, including nectar, pollen, food body, oil, resin, floral heat, shelter, and tryst sites (Simpson & Neff, 1981, 1983; Rands & Whitney, 2008). Some flowers offer floral tissues as a brood site reward for their pollinators (Sakai, 2002). Such nursery pollination (also brood-site pollination) systems are found in at least 15 families, including Agavaceae, Araceae, Arecaceae, Caryophyllaceae, Eupomatiaceae, Moraceae, Phyllanthaceae and Thymeleaceae (Sakai, 2002; Hossaert-McKey et al., 2010; Vislobokov et al., 2014; Zhang et al., 2021). Notably, several genera are pollinated by seed-predating insects, namely *Ficus* by agaonid wasps from the superfamily Chalcidoidea (Cruaud et al., 2012), *Yucca* by *Tegeticula* moths (Althoff et al., 2012), *Breynia*, *Glochidion* and *Phyllanthus* by *Epicephala* moths (Kawakita, 2010), and *Silene* by *Hadena* moths (Kephart et al, 2006). These flowers sacrifice some of their ovules as a food source for the pollinator broods, while other flowers offer other floral organs as the brood substrate for their pollinators, including perianth (*Eupomatia*; Armstrong & Irvine, 1990), resin-filled perianth (Schisandraceae; Luo et al., 2018), pollen grains (*Aspidistra xuansonensis*; Vislobokov et al., 2014), and male flower buds (*Phyllanthus*; Kawakita et al., 2022). These interactions can be highly specific and tightly co-evolved with their pollinator lineages. The brood-site flowers mediate interactions with their pollinators with various types of floral volatiles, primarily consisting of terpenoid, benzenoid or fatty acid derivative volatiles (Hossaert-McKey et al., 2010).

The combination of thinner cuticle layers, presence of floral rewards, and a ready supply of nutrients for future ovule development, renders flowers an excellent substrate for fungal growth (Ngugi & Scherm, 2006; Aleklett et al., 2014). In most scenarios, the fungi are either pathogenic or parasitic to the flowers (Ngugi & Scherm, 2006), and may reduce visitation and oviposition rate of the pollinators (Biere & Honders, 2006; Rering et al., 2021). Flower-fungus interactions can, however, be commensal or mutualistic (Rering et al., 2018; Vannette, 2020). Tripartite pollination systems involving fungal partners are perhaps most well-characterised in nectar yeast (Álvarez-Pérez et al., 2019). Nectar yeasts can serve as facultative mutualistic partners that can alter nectar nutrient value (Vannette et al., 2013) and odour (Yang et al., 2019), improving floral signalling and flower performance (Schaeffer et al., 2017), and may even mitigate the negative impact of herbivory (Deng et al., 2024). In most cases, the occurrence of nectar yeast seems to be stochastic among individuals in the populations, and permeate numerous flowering species in a community (de Vega et al., 2009; Herrera et al., 2009). In comparison, reports of tripartite pollination systems involving mutualistic filamentous fungal partners are much rarer in the literature, having so far been reported only in *Artocarpus integer* (Sakai et al., 2000) and *Artocarpus heterophyllus* (Gardner et al., 2018). These flowers attract gall midges (Cecidomyiinae, Diptera) as pollinators. The flowers of *Artocarpus heterophyllus* emit a cocktail of aliphatic esters, such as methyl 2-methylbutyrate, methyl tiglate and methyl isovalerate. The male inflorescences of these species are infested with *Choanephora* (*A. integer*) or *Rhizopus* fungi (*A. heterophyllus*), and the pollinating fungus gnats utilise these fungi as brood site rewards. Unlike nectar yeasts, the participation of the fungal partners is essential in these systems, and involves oviposition of the pollinators.

The pantropical family Annonaceae is largely pollinated by small beetles in the Nitidulidae, Curculionidae and Staphylinidae (Saunders, 2020). Members of the family almost always have the same floral Bauplan of one whorl of three sepals, two whorls of three petals, numerous spirally arranged stamens and an apocarpous gynoecium (van Heusden, 1992). The petals in most species form a more or less enclosed chamber in which pollinators consume food rewards and copulate. Most species have fleshy perianth parts that might provide adequate nutrients for pollinator broods, but assessment of nursery pollination is rare (Saunders, 2020). In particular, the Annonaceae genus *Meiogyne* has elaborate and thickened inner petal basal corrugation on its adaxial surface (van Heusden, 1992, 1994). Previous studies have established that this growth is rich in polysaccharides (Shao & Xu, 2015, as “*Oncodostigma hainanense*”; Xue, 2021). It has been suggested that in the Australian species *Meiogyne heteropetala*, inner petal corrugation might have provided tactile cues that encourage its beetle pollinators to lay eggs (Liu et al., 2024). Here, we present a pollination study of *Meiogyne hainanensis*, which is native to Hainan Island, China, and can grow up to 16 m tall. The inner petal corrugation of this species was reported to be prone to fungal infection (Shao & Xu, 2015). We use a combination of field observation, floral odour characterisation and metabarcoding to investigate the potential nursery pollination system of this species. There have been increasing research efforts to investigate floral fungus community. This study aims to provide insight into how flowers, insects and fungi achieve three-way mutualism.

## Materials and methods

### Study field sites

The tree species *Meiogyne hainanensis* (Annonaceae) (Fig. 1) was the focus of the current study. Field work was conducted during the peak flowering season (April–May) in Bawangling National Forest Park (BW), Hainan, China (two trees), Jianfengling National Forest Ecological Field Station (JF), Hainan, China (*ca.* 10 trees), and Huishan National Forest Park (HS), Hainan, China (*ca.* 10 trees) during 2018 and 2019, and a cultivated individual in South China Botanical Garden (SC), Guangdong, China in 2017. Individuals in BW and HS grew in tropical montane rainforests, while individuals in JS grew in a secondary tropical montane rainforest. The three field sites in Hainan were 50–150 km away from one another. The cumulative observation period lasted 71 days, consisting of *ca.* 380 hours.

**Fig. 1.**
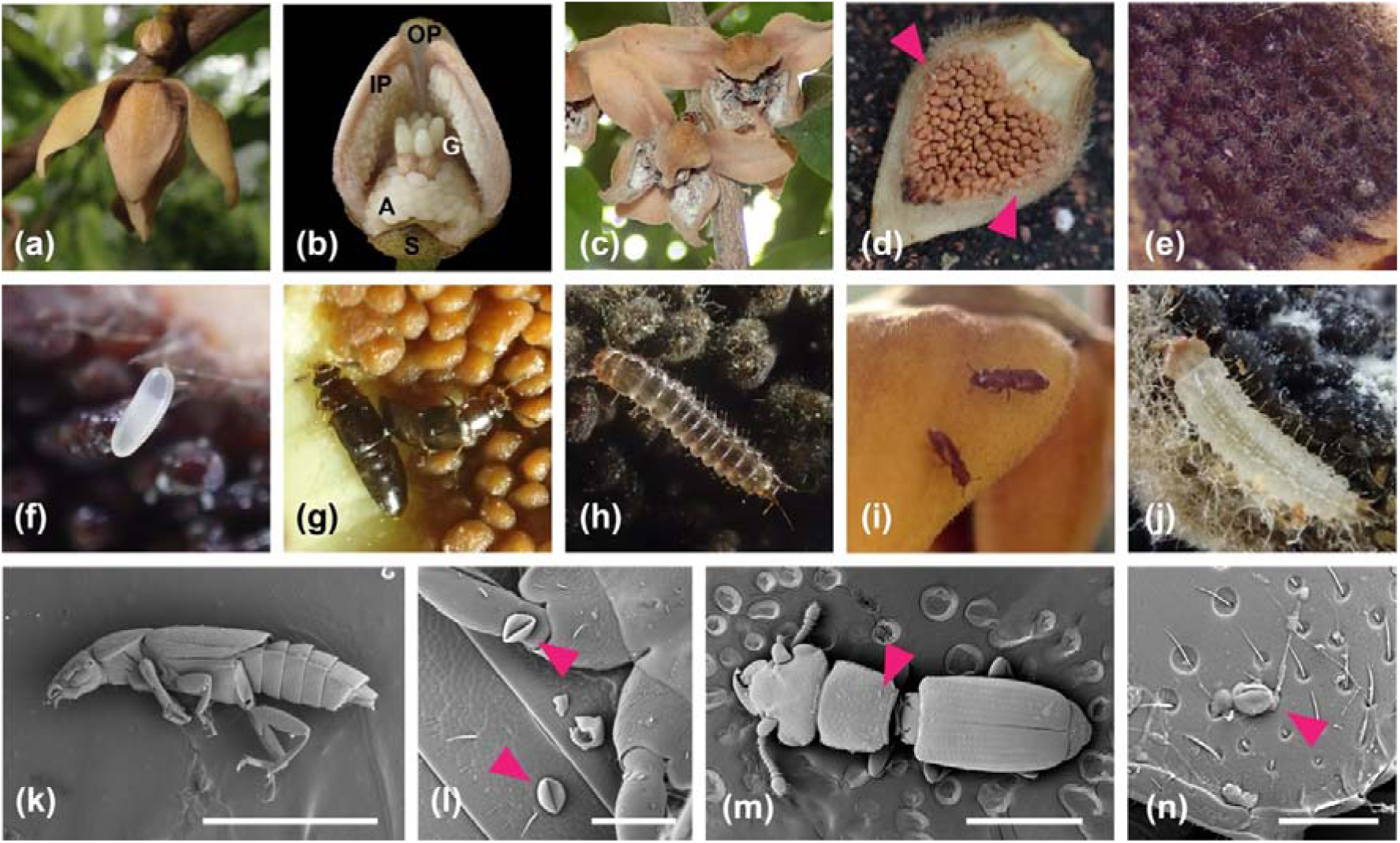
*Meiogyne hainanensis* and its pollinators. (a) pistillate-phase flower of *M. hainanensis*. (b) floral anatomy of early pistillate-phase flower (two outer petals and one inner petal removed). (c) post-anthetic flowers with prolific growth of filamentous fungi on the inner petal corrugation. (d) inner petal with corrugation (magenta arrows indicate beetle eggs deposited). (e) ascomycete growth on the inner petal of an early post-anthetic phase flower. (f) a beetle egg on the inner petal corrugation. (g) *Paraphloeostiba* sp. adults on the inner petal corrugation of overlap-phase flowers. (h) *Paraphloeostiba* sp. larva on the inner petal corrugation of a post-anthetic flower. (i) *Mimemodes* sp. adults on the outer petal of an overlap-phase flower. (j) *Mimemodes* sp. larva. (k–n) scanning electron micrographs of pistillate-phase visitors (magenta arrows indicate pollen deposition). (k–l) *Paraphloeostiba* sp. (m–n) *Mimemodes* sp. A: androecium; G: gynoecium; IP: inner petal; OP: outer petal; S: sepal. Scale bars: (k, m): 1 mm; (l, n): 0.1 mm.

### Floral ontogeny

To assess floral phenology, 50 flowers were tagged and hourly observations of perianth movement and receptivity of reproductive organs were made for ten individuals in BW, JS and SC. Because stigmatic peroxidase activity was detected as early as in the bud stage, stigma receptivity was instead defined as the duration between the secretion of stigmatic exudate and abscission of stigmas. The onset of staminate function was determined by dehiscence of anthers, and the end inferred by the breakdown of floral chamber. The pistillate phase is defined here as the period during which the flower exclusively has pistillate function; with the overlap phase defined as the period during which the flower has both pistillate and staminate function; and the staminate phase is defined as the period during which the flower has exclusively staminate function. A digital data-logger (Testo 176-T4, Testo, Titisee-Neustadt, Germany) connected to two type-K thermocouples (± 0.3°C accuracy) was used to assess floral temperature. One thermocouple was inserted into the floral chamber, and the other was placed 10 cm away from the flower to record ambient temperature. Temperatures were recorded every 10 min throughout anthesis.

### Floral visitors and Pollinators

Floral visitors often spend considerable time inside Annonaceae flowers (Gottsberger, 1988). Observations on floral visitor were made around the clock for the first 24 hours by carefully opening the floral chamber and inspecting floral visitors inside. Observation efforts were then reduced, with a 1-hour observation period conducted every 2 hours for the next week. Observation efforts were then concentrated towards the dawns and dusks, during which the inter-floral movement of the pollinators were most frequent. Pollinator observation was conducted in BW, JF and SC, but observation was logistically impractical in HS as the trees were too tall. Representative floral visitors were collected and either stored in 75% ethanol for identification, or killed in a -20°C freezer and silica-dried to assess presence of pollen grains. Because Annonaceae flowers are bisexual and protogynous, we assessed pollinators based on these criteria: (1) visitation frequency in relation to other floral visitors; (2) visitation to both sexual phases; and (3) presence of pollen grains on pistillate-phase floral visitors, which would demonstrate the inter-floral movement necessary to achieve pollination.

### Scent characterisation

The vegetative tissues, including leaves and flower buds of *Meiogyne hainanensis* produce a strong sesquiterpene-like smell to human olfaction. Composite sampling of Annonaceae flowers is recommended for the detection of volatiles at lower abundance (Goodrich et al., 2006), however. We therefore adopted the approach recommended by Goodrich et al. (2006) to include multiple flowers for each headspace sample. For each sample, three flowers were excised from the branch, and the excision points were sealed in parafilm and placed in a headspace bag (“Toppit” oven bag) to reduce wound-induced volatiles. To account for wound-associated volatiles, one flower bud was likewise excised and placed in a headspace bag without parafilm. In another setup, one undamaged anthetic flower attached on the plant was carefully bagged for sampling floral odour of non-wounded tissues. Post-anthetic flowers were invariably infested with fungi; fungus-associated volatiles were determined by comparison to sterilised post-anthetic flowers. Post-anthetic flowers were sterilised with 10% bleach solution for five consecutive days, each time for 5 min, then rinsed three times with Milli-Q ultrapure water. Six hours after the sterilisation on the last day, three sterilised post-anthetic flowers were likewise excised with the wound sealed with parafilm, and then transferred to a headspace bag. To assess the odour of mature fruits, intact fruits were removed from the trees and placed in headspace bags. To account for ambient contaminants, air controls were sampled every time floral scent was extracted, in which parafilm was placed in an empty headspace bag. Molecules detected in the air control were omitted in the floral samples. Samples were allowed to equilibriate in the headspace bags for 30 min. Floral volatiles were extracted using solid-phase microextraction (SPME) fibre with polydimethylsiloxane-divinylbenzene (PDMS-DVB; 65 mm film thick; Supelco, USA) coatings, with an adsorption period of 30 min. The SPME fibres were sealed in double-layered headspace bag during transportation to the laboratory at The University of Hong Kong. Analyses were performed within two days of extraction.

Gas-chromatography mass spectrometry (GC-MS) was performed in an Agilent 6890N/5973 gas chromatograph-mass selective detector. Volatiles were thermally desorbed at 250 °C for 1 min before being injected into a DB-WAX column (30 m × 0.25 mm, 0.25-mm-thick film; J&W). Flow was maintained at 1.5 ml/min with helium as the carrier gas. The column was kept at 50 °C for 5 min, followed by a ramp of 5 °C/min until reaching the final temperature at 230 °C, which was held for 5 min. Peaks were manually integrated and tentatively identified using the Wiley and National Institute of Standards and Technology mass spectral libraries (NIST, USA; 2020) with a threshold of 85% quality values. The compound identity was verified using either published standardised retention index values or by coinjecting with commercially available standards. Flower-specific volatiles were defined here as those that were found in at least intact flowers or composite flower samples, but not in flower bud samples. Wound-induced volatiles were defined as those shared between the flower bud and composite floral headspace, but not in intact flower samples. Vegetative background volatiles were defined as those shared among intact flower, composite flower, and flower bud samples.

### Fungal DNA extraction, amplification and sequencing

To identify the fungal composition on the post-anthetic flowers and the gut content of the pollinators, we performed ITS2 metabarcoding. Adult beetle pollinators from anthetic flowers and larvae on the inner petals of post-anthetic flowers were collected in 100% molecular-grade ethanol, with each sample composed of 10–20 individuals. Inner petals from post-anthetic flowers were collected in the field, stored in 50 ml sterile DNA-free centrifuge tubes and transported to laboratory in The University of Hong Kong within 6 h. Sample collection was performed with single-use sterile nitrile gloves. Samples were stored in a -80 °C freezer at the earliest opportunity. Laboratory bench and equipment were cleaned with 5% bleach solution before use. Prior to DNA extraction, beetle larvae and adults were rinsed in 1.5 ml 90% ethanol, 2% tween-80 solution thrice to remove fungal residue on the body surface. Since dissection and manipulation of the guts of small insects are often difficult and can be a source of contamination, surface-sterilised insects are frequently used as proxies for assessing the gut contents of small insects (Rassati et al., 2019; Veselská et al., 2023). DNA was extracted using Qiagen DNeasy Plant Mini Kit (Düsseldorf, Germany), following the manufacturer’s recommended protocol. Two sets of negative controls were performed: (1) DNA extraction was repeated with 100 µl UltraPure water (Invitrogen, USA) to control for potential kit contamination, and (2) the discarded disinfectant from the third rinsing round was collected and combined together based on insect species and developmental stage. DNA extraction were then repeated with 100 µl of the combined solutions. The ITS region is a standard DNA barcoding region for fungi, owing to its large barcode gap and higher amplification success rates (Schoch et al., 2012). The amplification of the full-length ITS often creates unnecessary chimera issues due to the conserved 5.8S region (Ihrmark et al., 2012), and therefore ITS2 regions were amplified using fungal universal primers (ITS3: 5’-GCATCGATGAAGAACGCAGC-3’; ITS4: 5’-TCCTCCGCTTATTGATATGC-3’; White et al., 1990) to characterise the fungal composition in the samples. While there are other more recently developed primers for the ITS2 regions with better performance (Ihrmark et al., 2012), the universal forward primer ITS3 was used here, because it contains a nucleotide mismatch to *M. hainanensis* ITS within 6 bp at the 3’ end, enabling more selective amplification of fungal DNA. The DNA samples were sent for commercial metabarcoding sequencing at Novogene. To construct the DNA libraries, the amplicon products were size-selected by 2% agarose gel electrophoresis, end-repaired, A-tailed, and ligated with adaptors. The libraries were sequenced on the Illumina NovaSeq 6000 platform to produce 250 bp paired-end raw reads. Primers were removed using Cutadapt (Martin, 2011). The data were filtered and denoised using DADA2 (Callahan et al., 2016). Reads less than 50 bp long or those containing Ns were removed, after which the pair-end filtered reads were merged. Chimeras were subsequently removed. The identities of amplicon sequence variants (ASVs) were annotated against the UNITE database using the “assigntaxonomy” function in DADA2 (Nilsson et al., 2019). Fungal ASVs were annotated by FUNGuildR (Nguyen et al., 2016). Rarefaction curve was constructed using the R package vegan (Oksanen et al., 2022) to assess if the sequencing depth was sufficient. All negative controls did not produce any visible bands after amplification, and therefore was not subjected to the subsequent library preparation or sequencing.

To assess similarity in fungal composition among samples, non-metric multidimensional scaling (NMDS) was performed. The compositions for fungal constituents were square-root transformed, and the Bray-Curtis dissimilarity index among the samples was calculated, which were then used to construct the NMDS plot with the R package vegan (Oksanen et al., 2022). The analysis of similarity (ANOSIM) was performed with 10,000 permutations to test if there is statistically significant difference in fungal communities among the groups.

## Results

### Floral phenology

The floral phenology of *Meiogyne hainanensis* is summarised in Fig. 2a. Floral odour was detected throughout day and night. The intensity was strongest to human olfaction around dusk and dawn, and was weakest around midnight. During the pistillate phase, the flower smelled fruity, with a hint of overripe fruit. During the staminate phase, the floral odour became more fermented to human olfaction. The flowers were not thermogenic. *Meiogyne hainanensis* produced bisexual flowers with a 2–3-day pistillate phase and *ca.* 9 h overlap phase. Due to logistic limitations, self-compatibility was not assessed. However, in the SC site, which consist of only one individual, no fruit set was observed for two consecutive years (2017–2018) despite flowering profusely, suggesting the plant is unlikely to be autogamous. The flower entered its pistillate phase during the first two days when production of stigmatic exudate and emission of scent was first initiated. However, the pollinators mostly arrived during dusk on day 2, and subsequently stayed in the flower for the rest of anthesis. The inner petals remained erect during most of anthesis, creating an enclosed floral chamber (Fig. 1a). Pollinators accessed the floral chamber via three basal apertures and one apical aperture. Anther dehiscence was first observed around 0900 h on day 4. Stamen abscission and spreading of inner petals were observed around 1500 h on day 4 within a very narrow period of about 1 h, effectively breaking down the floral chamber (Fig. 2a). The pollinators remained in the flower for another 2–4 h. Mass departure of pollinators was observed around dusk on the same day (day 4). This marked the end of anthesis, because of the breakdown of the floral chamber and pollinator departure, although stigma abscission was not observed until *ca.* 1000 h on day 5. During the period between the end of dusk on day 4 and stigma abscission on day 5, the flowers still emit a strong scent but pollinator occurrence was significantly reduced. The perianth, however, remained firmly attached to the receptacle at least until day 21. Retention of petals usually lasted up to two months. During this extensive post-anthetic phase, filamentous fungi were observed to grow extensively on the inner petal corrugations in all flowers (Fig. 1e). Visible fungal hyphae were observed as early as the first day of post-anthetic phase.

**Fig. 2.**
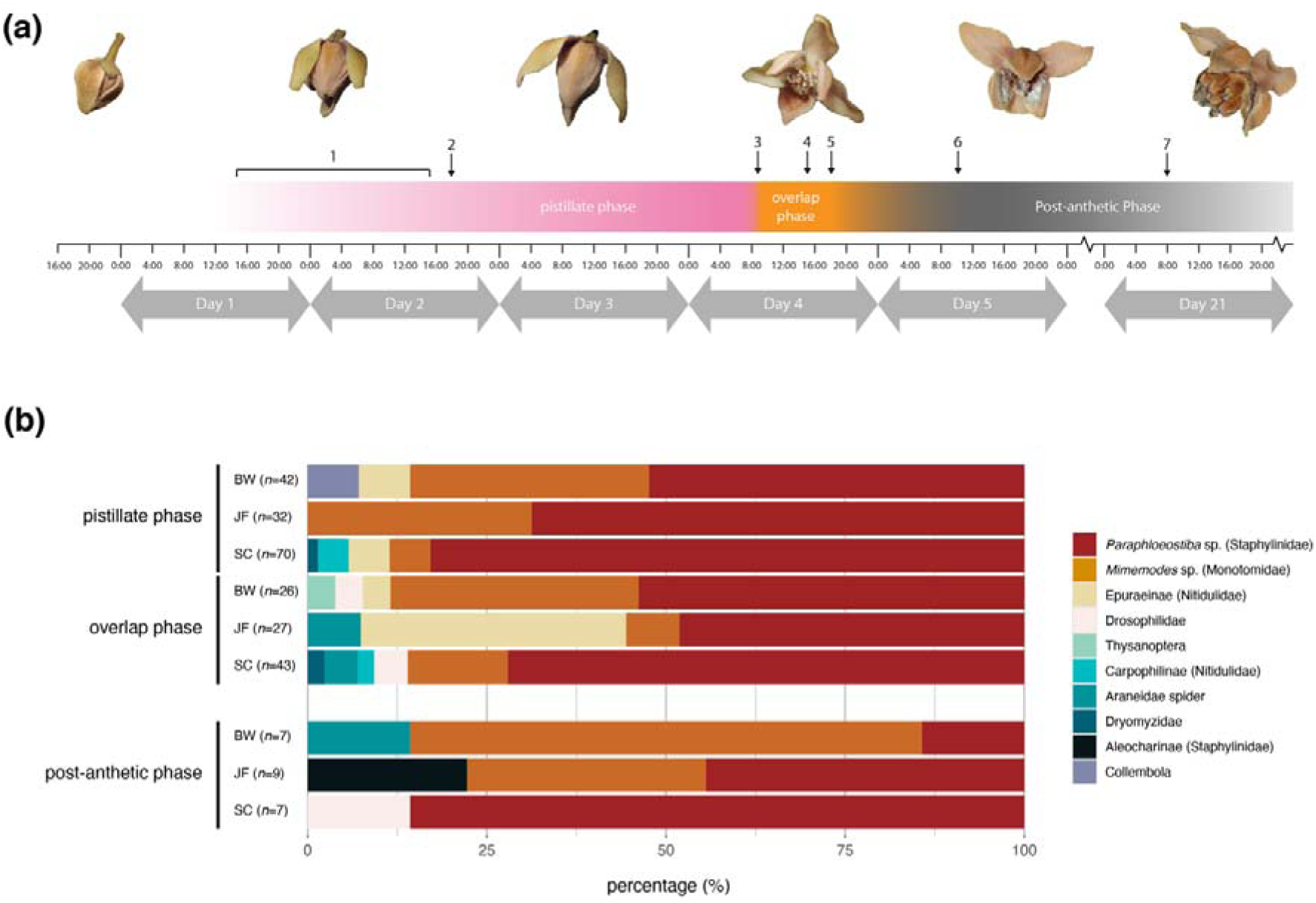
Floral phenology and floral visitors of *Meiogyne hainanensis*. (a) floral phenology of *Meiogyne hainanensis* (1: secretion of stigmatic exudate and emission of scent. 2: earliest arrival of beetle pollinator. 3: anther dehiscence. 4: stamen abscission and spreading of inner petals. 5: departure of beetle pollinators. 6: stigma abscission. 7: earliest abscission of inner petals). Inner petals usually remained attached for 1–2 months post-anthesis when the ovaries are developing. (b) floral visitors of *Meiogyne hainanensis*. *n* = the number of floral visitors. Field sites: BW: Bawangling; JF: Jianfengling; SC: South China Botanical Garden.

### Visitors and Pollinators

A total of 264 floral visitors were observed (Fig. 2b). The floral visitors comprised a Staphylinidae beetle *Paraphloeostiba* sp. (BW: 49.3%; JF:57.4%; SC: 79.2%), a Monotomidae beetle *Mimemodes* sp. (BW: 37.3%; JF:25.0%; SC:8.3%), a Nitidulidae beetle species from the subfamily Epuraeinae (BW: 16.7%; JF: 5.3%; SC: 3.3%), Drosophilidae flies (BW: 1.3%; JF: 0%; SC: 2.5%), a Staphylinidae beetle species from the subfamily Aleocharinae (BW: 4.0%), Nitidulidae beetles from the subfamily Carpophilinae (SC: 3.3%), Collembola (JF:3.3%), Dryomyzidae flies (BW: 1.7%), thrips (BW:1.3%), and Araneidae spiders (BW:1.3; JF: 1.7%; SC: 1.7%). Scanning electron microscopy revealed pollen deposition on the bodies of *Paraphloeostiba* sp. (family Staphylinidae) and *Mimemodes* sp. (family Monotomidae) retrieved from pistillate-phase flowers, suggesting they were likely to be effective pollinators (Fig. 1k–n). Both beetle species (Fig. 1g, i) were observed in both sexual phases, where they consumed stigmatic exudate and pollen grains and utilised the flower as a tryst site, and where they also gnawed on the inner petal corrugation. Mycangia were not observed in adult *Paraphloeostiba* sp. and *Mimemodes* sp. under SEM. Eggs were frequently observed on the inner petal corrugations during pistillate and overlap phases (Fig. 1e, f). Two species of beetle larvae (Fig. 1h, j) were frequently observed on inner petals of post-anthetic flowers (Fig. 1h, j). The larva as shown in Fig. 1j is morphologically congruent with the published description of Monotomidae (Jałoszyński, 2021), while the larva in Fig. 1h is morphologically highly similar to that of *Phloeonomus*, which is closely related to *Paraphloeostiba* (Staniec et al., 2016). Both species of larvae did not damage the developing ovaries nor other floral tissues, and were observed to exclusively graze on the fungal mycelia. Both *Paraphloeostiba* sp. and *Mimemodes* sp. larvae were able to grow and develop to late instar in the wild, sustaining themselves on the inner petal fungal hyphae. When the inner petals were transported to the laboratory, larvae of both species were likewise able to attain this size. Pupae were not observed on the inner petals of post-anthetic flowers in the wild, although *Mimemodes* sp. larvae were able to pupate and develop into adults in the laboratory. The larvae of *Paraphloeostiba* sp. were able to pupate but unable to reach adulthood, likely owing to suboptimal laboratory settings (25 °C; saturated humidity). The exact timing for egg hatching was hard to assess, since most eggs were well concealed under the crevices of the inner petal corrugations. It is thus difficult to assess the duration of larval stage, but generally most larvae for both species pupated 2–3 weeks after the end of anthesis.

### Floral scent composition

We detected a total of 70 volatiles from 11 classes across all organs (Table 1). In the anthetic flower samples, 30 volatiles were detected. In pistillate-phase flowers, 26 volatiles were detected, in which 56.43 ± 9.45% by relative abundance were flower-specific volatiles, 4.7 ± 1.94 % were wound-induced volatiles, and 38.88 ± 8.38% were vegetative volatiles that were present also in bud stage. In overlap-phase flowers, 27 volatiles were detected, in which 51.13 ± 9.77 % were flower-specific volatiles, 4.87 ± 2.09 % were wound-induced volatiles, and 44.00 ± 8.17 % were vegetative volatiles. While composite sampling created wound-induced compounds, this sampling strategy allowed the identification of low abundance floral molecules, accounting for 54.55% flower-specific chemical species in pistillate-phase flowers (9.92 ± 5.58 % by relative abundance), and 53.85% flower-specific chemical species in overlap-phase flowers (9.46 ± 2.39 % by relative abundance).

**Table 1.**
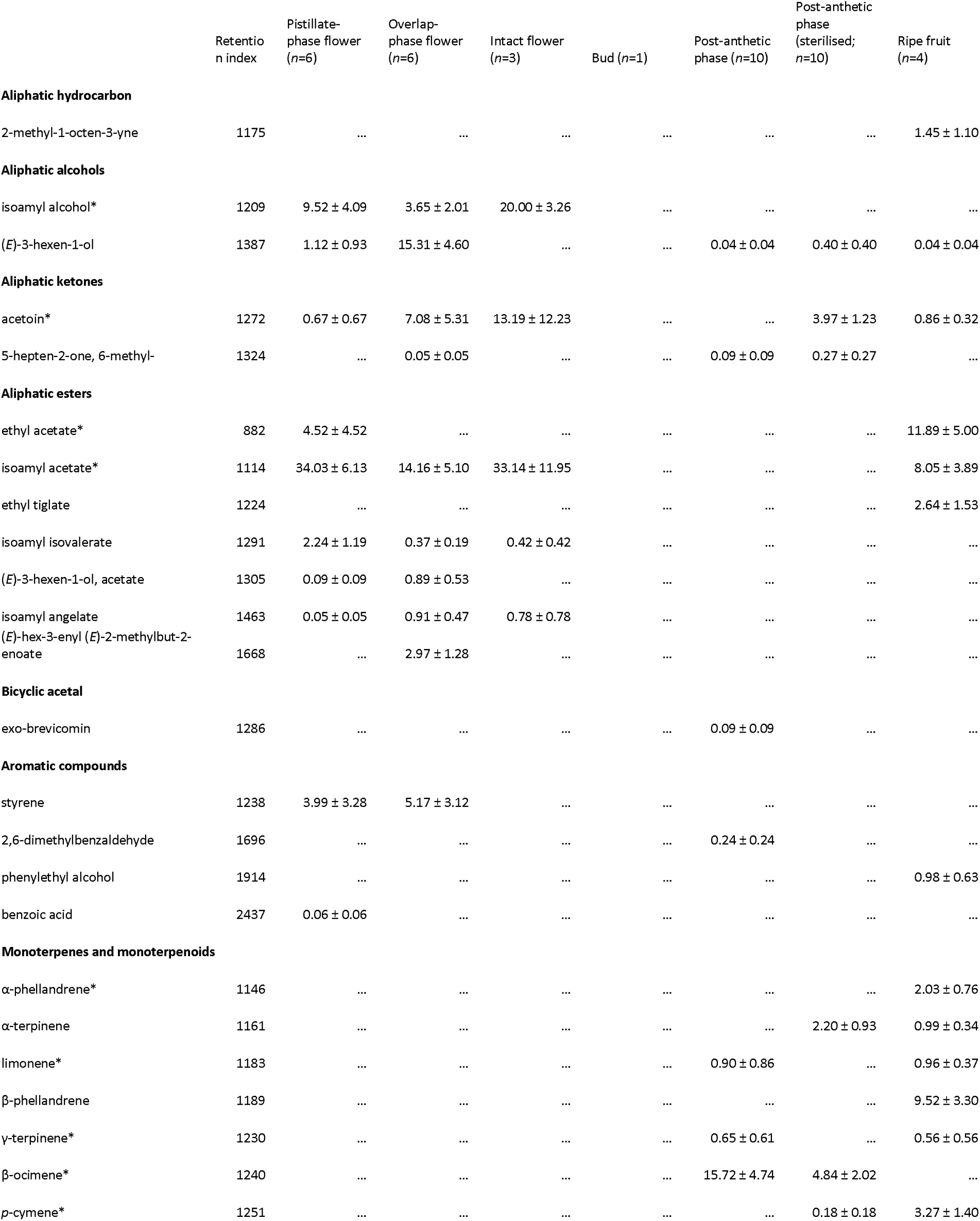
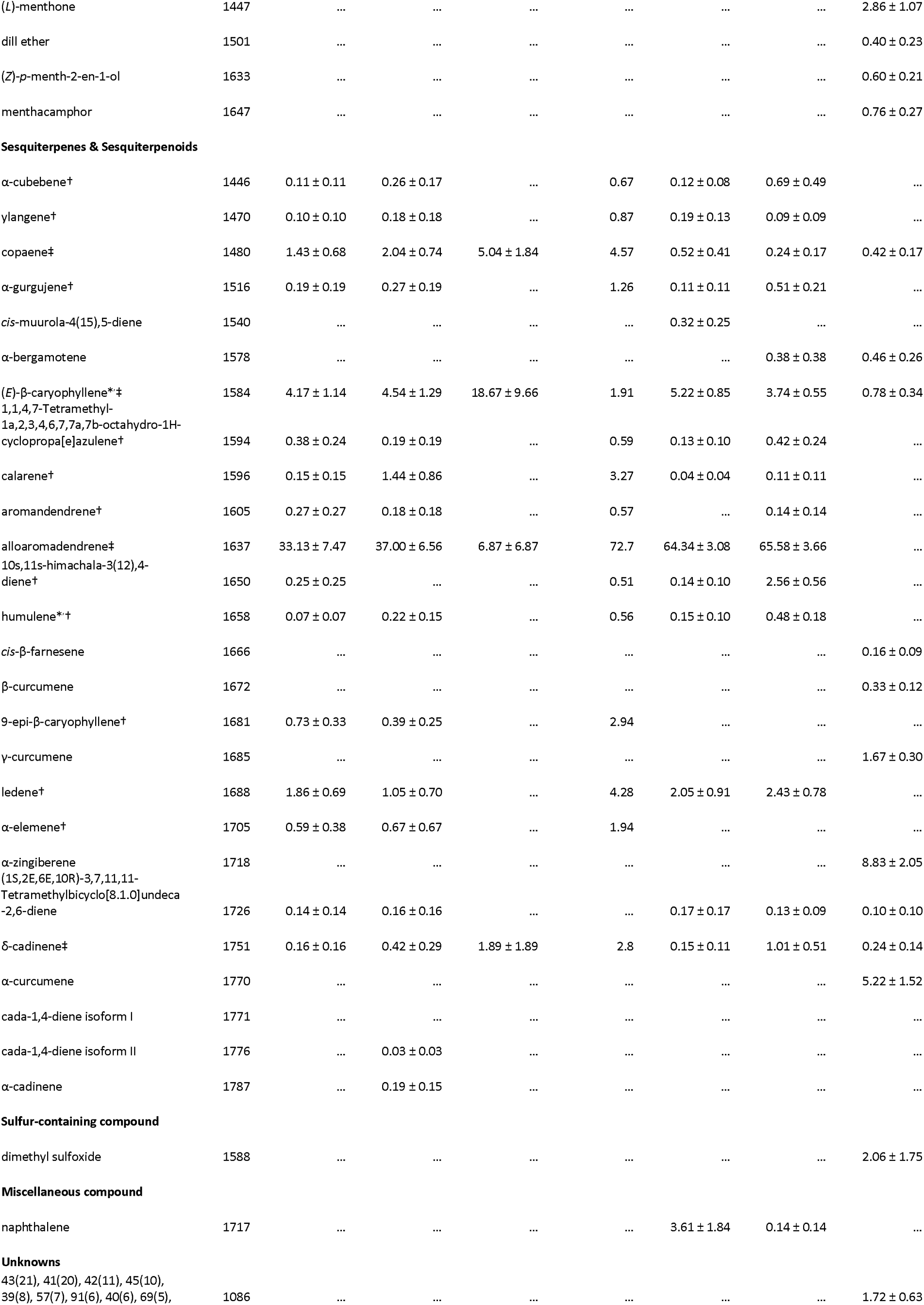
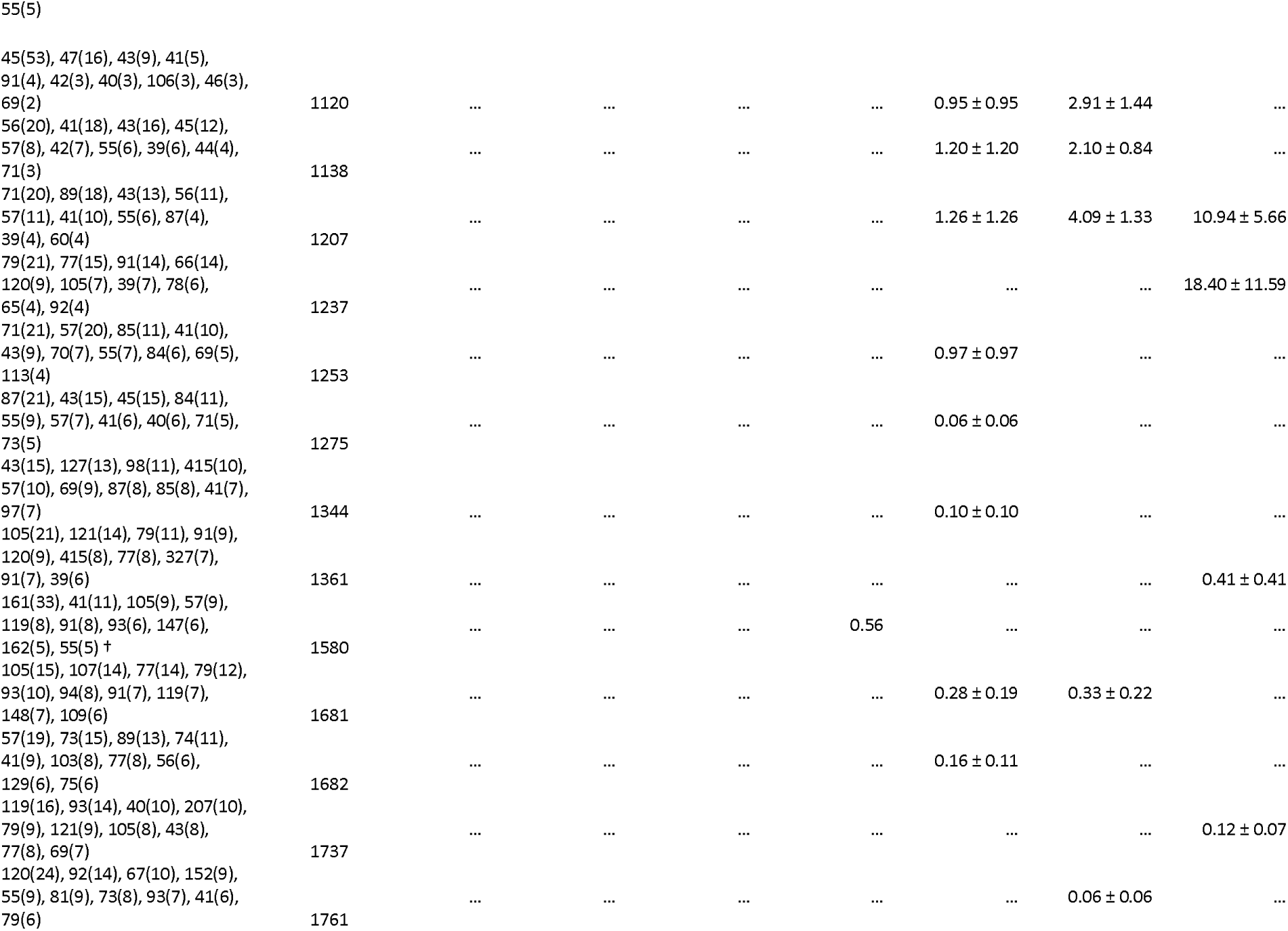
Volatile compounds of various organs of *Meiogyne hainanensis*. *: molecules verified by commercially available synthetic standards. Unknowns were represented by the top 10 most abundant ion fragments, followed by percentage abundance in parentheses. †: wound-induced volatile. ‡: vegetative background volatiles.

The most abundant floral volatile was the vegetative background volatile, alloaromadendrene (pistillate-phase: 33.13 ± 7.47%; overlap-phase: 37.00 ± 6.56%). Other major floral volatiles were mostly flower-specific compounds, including isoamyl acetate (pistillate-phase: 34.03 ± 6.13%, overlap-phase: 14.16 ± 5.10%), (*E*)-3-Hexen-1-ol (pistillate-phase: 1.12 ± 0.93%, overlap-phase: 15.31 ± 4.60 %), isoamyl alcohol (pistillate-phase: 9.52 ± 4.09%, overlap-phase: 3.65 ± 2.01%), acetoin (pistillate-phase: 0.67 ± 0.67%, overlap-phase: 7.08 ± 5.31%), styrene (pistillate-phase: 3.99 ± 3.28%, overlap-phase: 5.17 ± 3.12%), and ethyl acetate (4.52 ± 4.52%).

The headspace of post-anthetic flowers mostly consisted of sesquiterpenes (73.72 ± 4.15%), most of which were vegetative background volatiles that were also found in flower buds, including alloaromadendrene (64.34 ± 3.08%). A total of 26.28 ± 4.15% volatiles of post-anthetic flowers were not detected in anthetic flowers, composed of primarily beta-ocimene (15.72 ± 4.74%) and naphthalene (3.61 ± 1.84%). Sterilised post-anthetic flowers produced a largely similar odour profile, in which 96.83 ± 1.19% of the volatiles was also present in the non-sterile post-anthetic flowers. This suggests that most of the volatiles of post-anthetic flowers originated from the plant itself, and there was a limited influence of microbes on the odour of post-anthetic flowers. Some molecules, however, were only emitted after sterilisation of the inner petal corrugation, most notably acetoin (3.97 ± 1.23%) and α-terpinene (2.20 ± 0.93%). These were likely artefact emission of plant volatiles created by weakened plant cell walls and cuticles under disinfectants.

The mature fruit odour was largely dominated by two unknown molecules (18.40 ± 11.59%; 10.94 ± 5.66%), two aliphatic esters, namely ethyl acetate (11.89 ± 5.00%) and isoamyl acetate (8.05 ± 3.89%), and the sesquiterpene, α-Zingiberene (8.83 ± 2.05%). Both ethyl acetate and isoamyl acetate were also detected in the floral headspaces.

### Metabarcoding

The ITS2 amplicon metabarcoding sequencing generated 5,529,588 pair-end reads in total. After filtering and denoising to remove singletons, chimeras and reads with Ns, 4,283,145 pair-end reads were recovered, with a recovery rate of 77.6 ± 1.3%. The average amplicon length is 324 bp (67–432 bp). The rarefaction curves suggest that the sequencing depth was sufficient for reliable estimate of species richness in all fungal biomes (Fig. 3a). A total of 482 ASVs were recovered. On average, 99.7 ± 8.84 ASVs were detected in the inner petal corrugation; 179.4 ± 21.5 ASVs were detected in *Paraphloeostiba* sp. larvae; 209.8 ± 63.6 ASVs were detected in *Paraphloeostiba* sp. adults; 225 ± 32.2 ASVs were detected in *Mimemodes* sp. larvae; and 215 ASVs were detected in *Mimemodes* sp. adults. In total, the most frequently recovered fungal ASV belongs to *Fusarium waltergamsii* ASV1 (1,743,065 reads), *Penicillium paxilli* (261,529 reads), *Fusarium* sp. (253,818 reads), Fungi incertae sedi (230,078 reads), and *Cladosporium* sp. (184,557 reads). The fungal microbiome of post-anthetic inner petal corrugation was largely similar across all field sites, composed of mostly ascomycete fungi, including *Fusarium waltergamsii*, *Penicillium paxilli* and *Cladosporium* species. The major fungal taxa in *Paraphloeostiba* sp. and *Mimemodes* sp. larvae were similar to that of the inner petal corrugation. FUNGuildR yielded largely ambiguous results which are not presented here. ANOVA analysis detected a marginal statistical significance of Shannon index among groups (degree of freedom=4; sum of square=2.516; *F*-value=3.17; *p*=0.029*). However, after taking multiple comparison into account, the post-hoc Tukey’s HSD test failed to detect any pair-wise significant difference.

**Fig. 3.**
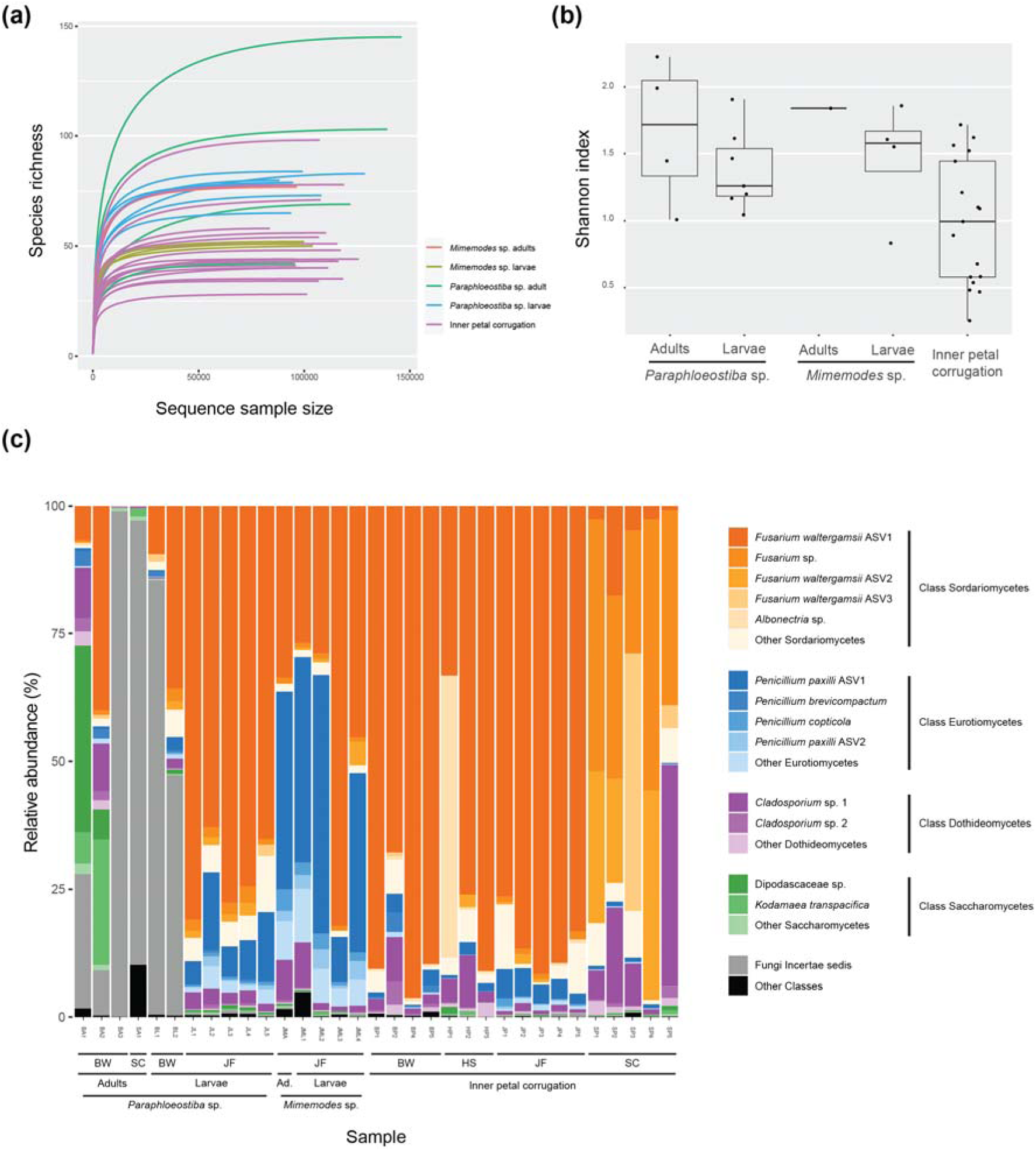
Fungal biome of a total of 33 samples of inner petals of *Meiogyne hainanensis* and the guts of adults and larvae of *Paraphloeostiba* sp. and *Mimemodes* sp. based on ITS2 metabarcoding sequencing. (a) Rarefaction curve showing sufficient sequencing depth. (b) Shannon index showing the alpha diversity of the 33 samples grouped by sample origin. (c) Composition of the ASVs detected in ITS2 metabarcoding sequencing. Sordariomycetes, Eurotiomycetes, Dothideomycetes and Saccharomycetes are classes that belong to the Ascomycota. Ad.: adult. Field sites: BW: Bawangling; HS: Huishan; JF: Jianfengling; SC: South China Botanical Garden.

Non-metric multidimensional scaling (NMDS) plot of the fungal biomes of the inner petal corrugations and the gut contents of *Paraphloeostiba* sp. and *Mimemodes* sp. adult and larvae is illustrated in Fig. 4 (2D stress value=0.15). The global ANOSIM analysis detected a significant difference among groups (*R*=0.23, *p=*0.0240*). Post-hoc pairwise ANOSIM revealed that there was no significant differences between the inner petal corrugation and *Mimemodes* sp. adult gut contents (*p*=0.340; *p_adj._*=0.396), nor with *Mimemodes* sp. larva gut contents (*p*=0.317; *p_adj._*=0.396). Likewise, there was no difference between *Mimemodes* sp. adult and larva gut contents (*p*=0.4; *p_adj._*=0.4). These findings suggest that both *Mimemodes* sp. adult and larvae consume fungal substrate highly similar to those on the inner petal corrugation. For *Paraphloeostiba* sp., there was no significant difference between larva gut contents and inner petal corrugation (*p.*=0.273; *p_adj._*=0.396), suggesting that larvae consumed the fungal community on inner petal corrugation. However, the fungal ITS2 composition of *Paraphloeostiba* sp. adult gut contents and inner petal corrugation was significantly different (*p*=0.001***; *p_adj._*=0.009**). While the *Fusarium*, *Penicillium* and *Cladosporium* ASVs were present, the fungal compositions of the guts of *Paraphloeostiba* adults and larvae were also significantly distinct from one another (*p*=0.006**; *p_adj._*=0.014*), suggesting a partitioning in diet between *Paraphloeostiba* sp. adults and larvae.

**Fig. 4.**
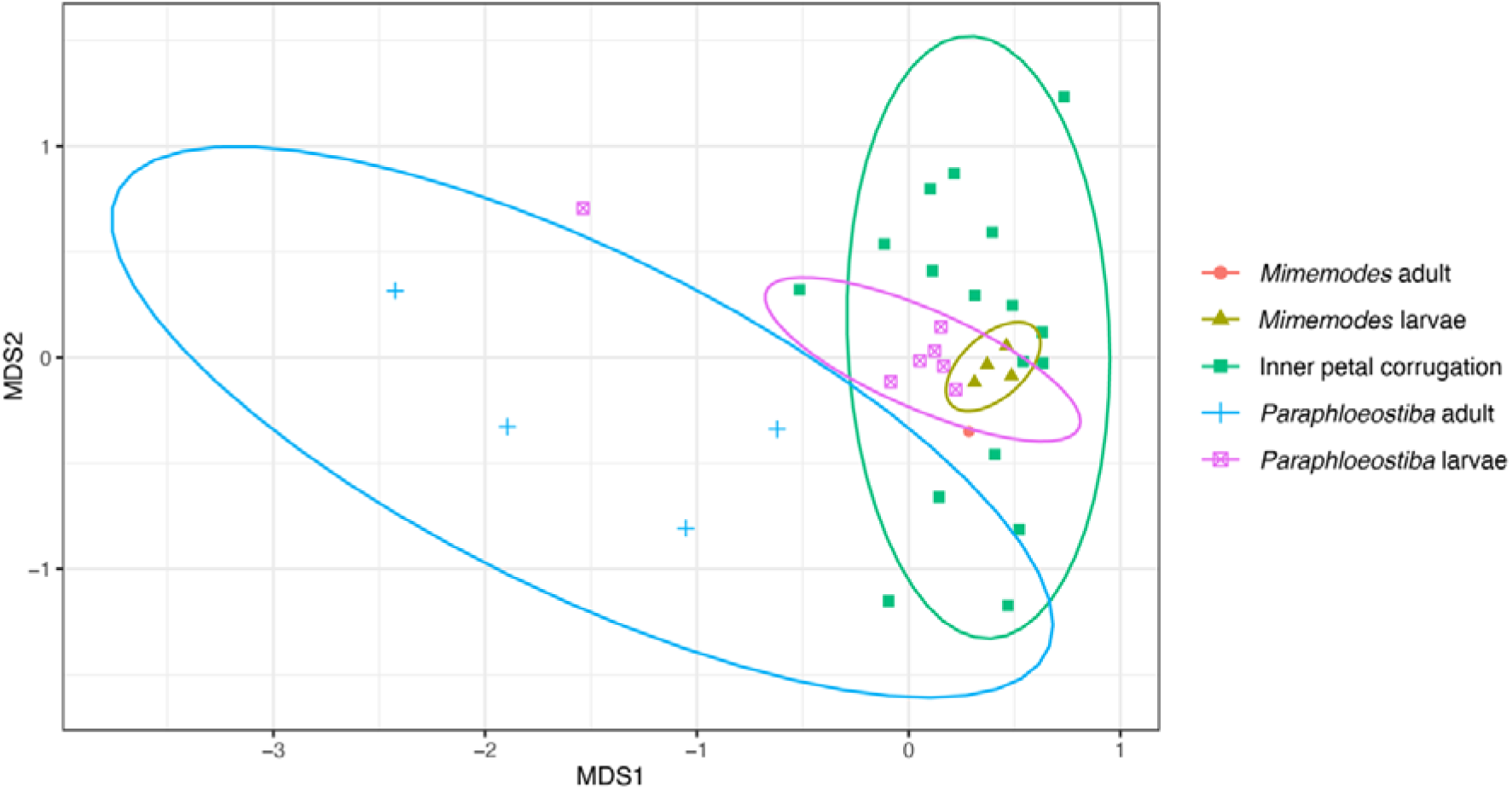
Non-metric multidimensional scaling plot of the fungal composition of inner petal corrugations of *Meiogyne hainanensis* and gut fungal composition of *Mimemodes* sp. and *Paraphloeostiba* sp. adults and larvae inferred by fungal ITS2 metabarcoding with 90% confidence interval ellipses. 2D stress level = 0.153; ANOSIM global *R*= 0.23; *p=*0.024*.

## Discussion

The findings of the current study suggest that *Meiogyne hainanensis* has a tripartite nursery pollination system (Fig. 5), involving two pollinators, *Paraphloeostiba* sp. and *Mimemodes* sp. (Fig. 1), and multiple ascomycete fungal partners, including *Fusarium* spp., *Penicillium* spp. and *Cladosporium* spp. (Fig. 3c). In this system, the beetles *Paraphloeostiba* sp. and *Mimemodes* sp. pollinated the flowers and consumed floral rewards, including pollen, stigmatic exudate and food body in the form of inner petal corrugation. The two beetle species also oviposited onto the inner petal corrugation. After anthesis, abscission of the petals of *M. hainanensis* was delayed and the inner petal corrugation served as a growth substrate for ascomycete fungi. The fungal mycelia were consumed by the larvae of *Paraphloeostiba* sp. and *Mimemodes* sp. Larvae of both species were able to attain the size of the late instars *in situ*, suggesting that *M. hainanensis* offers an honest brood site reward.

**Fig. 5.**
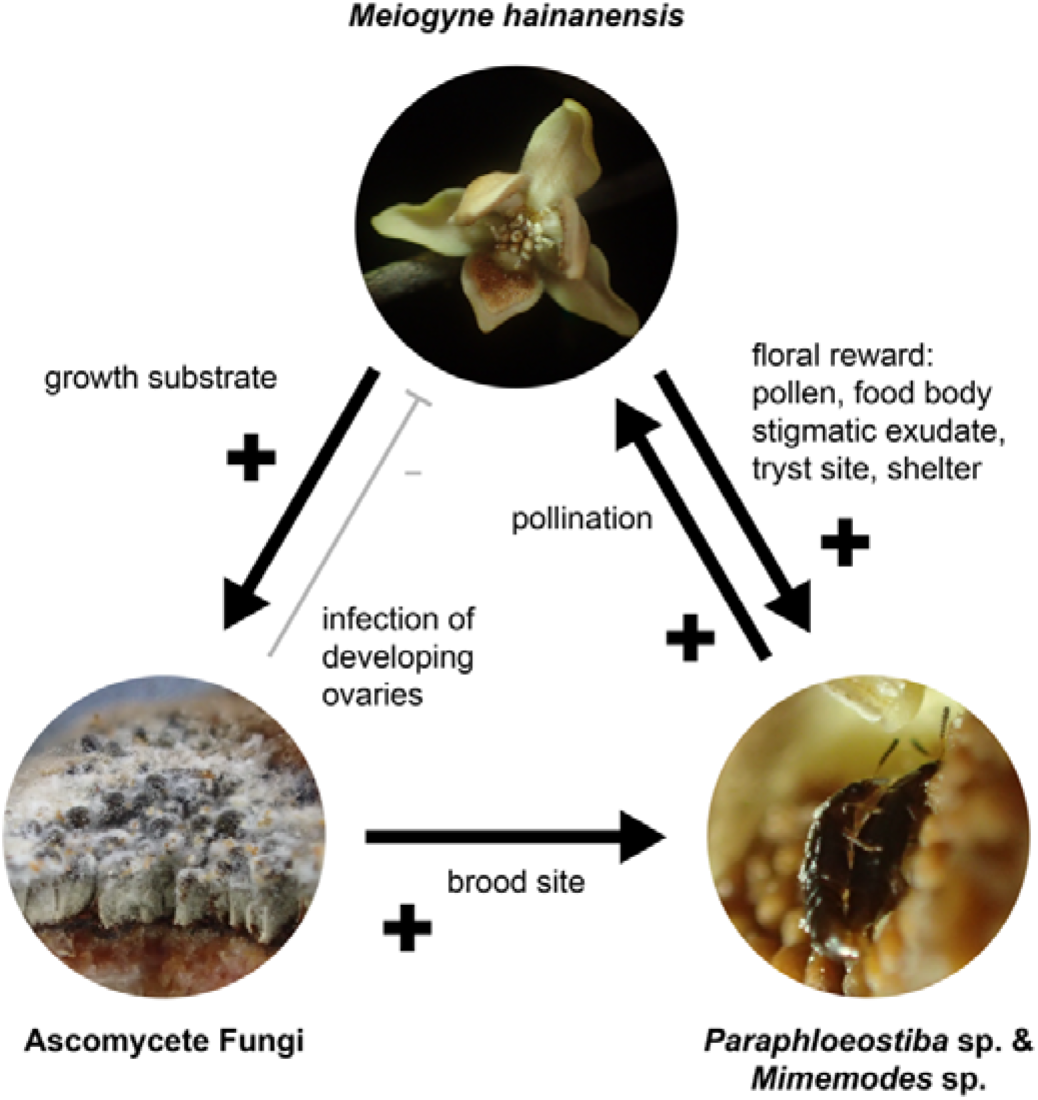
Tripartite interactions among *M. hainanensis*, ascomycete fungi and *Paraphloeostiba* sp. and *Mimemodes* pollinators. Arrows indicate the direction of the interactions from the givers to the receivers. Plus signs indicate positive interactions, while a minus sign signifies a negative interaction.

### Life history of the pollinators

The two pollinators, *Paraphloeostiba* sp. and *Mimemodes* sp. represented the majority of the floral visitors (Fig. 2b). Members of these two genera, *Paraphloeostiba* and *Mimemodes* have previously been reported to visit *Annona* flowers (Annonaceae) (Thayer et al., 2003; Higuchi et al., 2014). Utilisation of monotomid beetles as pollinators including *Mimemodes* is rare in Annonaceae, and has only been reported in *Annona cherimola* and its hybrid (Higuchi et al., 2014; Jenkins et al., 2015). Staphylinidae-pollinated lineages in Annonaceae, however, is much more common, including the genera *Annona*, *Anaxagorea*, and *Xylopia* (Jürgens et al., 2000). The pollinator genus *Paraphloeostiba* was reported to associate with fermenting fruits (Kuschel, 1990; Thayer et al., 2003) and bamboo-rotting fungi (Shavrin, 2017). The other pollinator genus *Mimemodes* is associated with decomposing wood (Grove, 2002), and has been reported to be an egg predator of a pine bark beetle residing in their fungus gallery (Kishi, 1970), or utilising fungal smut disease galls (Yoshitomi, 2017). Available documentation of the natural history of the pollinator genera indicates a close association with fungi, largely aligning with our observation of mycetophagous behaviour of the larvae, as well as the fungal constituents of their gut contents.

In the current study, both *Paraphloeostiba* sp. and *Mimemodes* sp. utilise the fungi growing on the inner petal corrugations of *M. hainanensis* as the larval food source, on which they develop into late instar. The larval stage lasts *ca.* 2–3 weeks, well under the typical duration in which the petal remained attached to the plant (1–2 months). They, however, do not pupate in the flowers. In their review on the behaviour of immature insects, Hagstrum and Subramanyam (2010) listed 59 families of insects which depart their larval feeding sites to search for pupation venues (typically soil or leaf litter), suggested partitioning of brood site and pupation site is common in insects. It was suggested that by departing their feeding sites, pupation can take place in more favourable microclimate with reduced predation risks (McPheron & Broce, 1996). While the beetles did not complete their life cycle on *M. hainanensis,* the flowers offer food source for beetle adults and larvae and tryst site for the adult beetles.

### Floral odour and fruity aromas

The vegetative molecules, especially the sesquiterpenes, constituted a significant part of the floral headspace of *Meiogyne hainanensis* (Table 1). Sesquiterpenes are volatiles commonly present in the essential oils of Annonaceae vegetative tissues (Fournier et al., 1999) and, in general, were reported as deterrents against antagonistic insects or suppressors of pathogenic attacks (Cheng et al., 2007). Flower-specific molecules included isoamyl acetate, isoamyl isovalerate, acetoin and isoamyl alcohol (Table 1). In particular, isoamyl acetate, isoamyl isovalerate and isoamyl alcohol have been reported as aroma molecules of various fruits, including banana (Zhu et al., 2018), jackfruit (Swords et al., 1978; Maia et al., 2004) and cherimoya (Ferreira et al., 2009). Acetoin, on the other hand, is a volatile associated with fermentation (Romano & Suzzi, 1996). Together, these volatiles may explain why the flowers smell like overripe fruits to human perception. Post-anthetic flower sterilisation experiments suggest that the fungi themselves had very little influence on the odour of post-anthetic flowers. This suggests that instead of directly exploiting cues offered by the fungi, the anthetic flowers probably only exploit the signal of a substrate that is likely associated with fungal growth, i.e. scent of overripe fruits. Indeed, fungal growth was almost always present on the inner petal corrugation of post-anthetic flowers. The flowers therefore provide an honest signal of brood site to the pollinators.

Interestingly, as the flower progressed from pistillate to overlap phase, the floral odour became incrementally more “rotten” to human olfaction, presumably corresponding to the higher level of acetoin in overlap-phase flowers (Table 1). Dimorphism of olfactory cues between sexual phases is not unusual in Annonaceae (Goodrich & Raguso, 2009). Progressive changes in floral scent to more rotten fruity odour during staminate/overlap phase have also been reported in the congeneric *Meiogyne virgata* (Silberbauer-Gottsberger et al., 2003). Because male fitness is limited by the ability of flowers to export pollen while female fitness is limited by the resources that can be invested to produce diaspores, the theory of sex allocation predicts that the staminate-phase flowers (in this case, overlap-phase) would evolve to be more attractive to pollinators (Stanton & Preston, 1988). Tsukada et al. (2017), however, suggested that *Annona* flowers might have instead employed a push-pull-like strategy (see also Terry et al., 2007), in which staminate-phase flower is less attractive than pistillate-phase flower, for encouraging pollinator movement from staminate-phase to pistillate-phase flower. It is unclear how sexual dimorphism in floral odour would affect pollination in *M. hainanensis* and the wider Annonaceae.

### Function of the inner petal corrugation

The basal inner petal corrugation is one of the two diagnostic characters for members of the genus *Meiogyne*, alongside the elongated connective of the innermost stamen whorl (van Heusden, 1994). This inner petal growth was previously interpreted as a gland (van Heusden, 1992, 1994), although this structure is rich in polysaccharide and devoid of any secretory ducts or openings (Shao & Xu, 2015; Xue et al., 2021), unlike other nectariferous Annonaceae species (Silberbauer-Gottsberger et al., 2003). Our study has identified that the inner petal corrugation in *M. hainanensis* offers food rewards for adult pollinators and brood substrate for pollinator larvae indirectly via its fungal partners. This observation aligns with a previous study that showed the corrugation in *M. hainanensis* was prone to fungal infection (Shao & Xu, 2015, as “*Oncodostigma hainanense*”). In contrast, the inner petal corrugation of the congeneric aerial litter mimic *Meiogyne heteropetala* appeared to provide tactile cues for oviposition (Liu et al., 2024). For *M. heteropetala*, although the inner petal corrugation was able to sustain the insect larvae to adulthood in the laboratory, the larvae mostly perished *in situ*, likely because the forest floor was unable to provide optimal environment for the insect broods on fallen petals. A combination of sufficient nutrients and optimal microhabitats was likely required for offering authentic nursery reward. This may additionally explain why *M. hainanensis* in the current study displayed highly unusual floral phenology, in which petals were retained on the receptacle for 1–2 months after anthesis (Fig. 2a): the arboreal environment might have provided crucial microhabitats for brood development, though this remains to be tested in *M. hainanensis*. In general, although pollinator copulation is commonly observed in Annonaceae flowers, evidence for nursery pollination is contentious and scarce in the family (Saunders, 2020). More detailed studies are needed to assess how widespread nursery pollination systems are in the Annonaceae.

### Tripartite pollination with fungal partners

While facultative tripartite pollination systems involving nectar yeast and other nectar microbes are common (de Vega et al., 2009; Herrera et al., 2009; Vannette, 2020), tripartite pollination systems involving essential filamentous fungal partners are much rarer. Such systems have been reported in *Artocarpus* (Moraceae), specifically *A. heterophyllus* and *A. integer*, in which gall midge pollinators oviposit in male inflorescences infested by Mucoralean fungi (McMillan, 1974, 1986; Sakai et al., 2000; Gardner et al., 2018). *Meiogyne hainanensis* flowers utilise a community of ascomycete fungi, including *Fusarium* spp., *Penicillium* spp. and *Cladosporium* spp. Most fungal partners of *M. hainanensis* were filamentous fungi. These genera are cosmopolitan saprophytes and plant pathogens (Zhang et al., 2006; Bensch et al., 2012; Visagie, 2016), and are commonly associated with rots or wilts, including rots in pre-and post-harvest fruits (McMillan, 1974, 1986; Sanderson & Spotts, 1995; Briceño & Latorre, 2008; Bakar et al., 2013; Visagie, 2016; Zakaria, 2023). *Fusarium*, *Penicillium* and *Cladosporium* species are responsible for the rotting of ripe fruits in *Annona* (Cambero-Ayón et al., 2019; González-Ruíz et al., 2021), with *Fusarium* and *Penicillium* spp. additionally pathogenic to mature fruits of *Asimina* (Ivan et al., 2020). The most abundant fungal partner *Fusarium waltergamsii* has been found on *Ficus* fruits (Chaverri & Chaverri, 2022), while *Penicillium paxillii* and a *Fusarium* sp. were previously isolated on the epicarps of multiple *Citrus* species (Moreno Rueda et al., 2013). There is a surprisingly low variation in microbe composition on the inner petal corrugation across populations of *M. hainanensis*. Floral microbes in other systems are usually strongly controlled by physiological filters, such as osmotic pressure and antifungal metabolites (Herrera & Pozo, 2010). In a study that attempt to characterise the molecular basis for such microbial filter, it was found that around 21% of nectar cDNAs are defence-related (Thornburg et al., 2003). Characterisation of transcriptome and metabolome of the inner petal corrugation of *M. hainanensis* may offer information on how the relationship with the fungal partners is maintained biochemically.

The *M. hainanensis* system is mutualistic as a whole, in which the plant gains pollination service, and the pollinating beetles gain floral rewards in the form of food source and an oviposition site, while the fungi utilise the inner petal corrugation as a growth substrate. When dissecting this three-way interaction and assessing its individual interactions, the flower-insect interaction is mutualistic but the nature of fungus-flower interaction and fungus-insect interaction may be ambiguous. The ascomycete fungi are probably largely commensal and occasionally pathogenic to the plant. Field observations suggest that fungal mycelia were largely confined to the inner petal corrugations. However, since the inner petals were retained after anthesis, early developing ovaries are in close proximity to the fungus-infested petal corrugation. Mycelia were sometimes seen spreading to infect developing ovaries during the wet year of 2019, although not in 2017 and 2018. Nevertheless, such negative influence on ovule development is likely much less severe in comparison to nursery systems that utilise seed-predating pollinators, in which the plant can face up to around 50% seed reduction (Kephart et al., 2006). It is unclear whether the adult beetles serve as vectors for fungal dispersal. Outside of the flowers, the beetle pollinators probably feed on ascomycete fungi on other substrates. For instance, *Paraphloeostiba* are known to associate with rotten fruits (Kuschel, 1990; Thayer et al., 2003). Both beetles lack specialised structures for dispersing fungal spores, i.e. mycangia. *Fusarium*, *Penicillium* and *Cladosporium* can be dispersed through airborne innocula (Rodríguez-Rajo et al., 2005; Halstensen et al., 2006; O’Gorman & Fuller, 2008); it is therefore unclear whether the pollinators effectively contribute to fungal spore dispersal. Characterisation of fungal biomes in pollinator-excluded post-anthetic flowers may shed light on the role of the pollinators as spore dispersers.

Compared to other nursery pollination in which the pollinator larvae directly feed on perianth tissue, such as Schisandraceae (Luo et al., 2018) and *Eupomatia* (Armstrong & Irvine, 1990), the participation of fungal partners does not seem to confer additional reproductive benefits. Apart from additional risks of fungal attack to developing ovaries, the fungal tripartite nursery pollination system in *M. hainanensis* is also associated with energy wastage due to the increased number of trophic levels involved and potentially prolonged maintenance of petal tissues. Beetle larvae can consume around 48% perianth in dry mass in the bipartite nursery flower *Eupomatia* (Armstrong & Irvine, 1990). The staggering extent of resources required may explain why *M. hainanensis* evolved highly thickened and fleshy inner petals (Fig. 1b, 1d). These additional metabolic costs may limit the energy that can be invested to fruit and seed development, potentially reducing the female fitness of the plant (Stanton & Preston, 1988).

### Influence of fungal biome on olfactory signalling to pollinators

Apart from offering a food source and changing the nutrient content of floral rewards (Vannette & Fukami, 2018), the floral microbiome may also directly alter floral odour (Golonka et al., 2014; Rering et al., 2018), and in another Annonaceae lineage, *Asimina triloba*, it enhances the olfactory attraction of pollinators (Martin, 2021). For *M. hainanensis*, however, fungal growth was not observed until after anthesis, and is therefore unlikely to directly contribute significantly to the olfactory signalling of anthetic flowers. Nonetheless, towards the peak and the end of the flowering season, fungus-infested post-anthetic flowers are accumulate to an appreciable number on branches. The fungal ASVs of the inner petal corrugation also constituted a part of the diet of adult *Mimemodes* sp. (Fig. 3c). Thus, post-anthetic flowers may exert a magnet effect (Laverty, 1992), in which a third-party rewarding substrate locally increases the abundance of pollinators, thereby creating a spill-over effect that indirectly facilitates pollination success of the co-occurring focal flower: in the case of *M. hainanensis*, fungi in post-anthetic flowers may potentially act as a magnet and indirectly promote floral visitation. While this is a plausible hypothesis, we detected only a low level of fungus-associated volatiles in post-anthetic flowers (Table 1). More importantly, considerably more adult pollinators were observed in the anthetic flowers than in post-anthetic flowers (Fig. 2b). Both observations do not seem to support the postulation of magnet effect. In contrast to nectar yeast (Golonka et al., 2014; Martin, 2021), the fungal biome in post-anthetic flowers of *M. hainanensis* is unlikely to influence the olfactory signalling to pollinators.

### Possible origin of tripartite nursery pollination systems

There appears to be a convergence towards emission of fruity odours in tripartite nursery flowers. The non-vegetative floral volatiles ethyl acetate, isoamyl acetate and acetoin were also found in fruit headspaces of *M. hainanensis* (Table 1). Similarly, in another tripartite nursery flower species *Artocarpus heterophyllus*, the flowers and fruits share identical aliphatic esters and alcohol, including methyl isovalerate, 2-methylbutyl acetate, pentyl acetate, hexyl acetate and 2-methyl-1-butanol (Gardner et al., 2018). Aliphatic esters are common aromas emitted by ripe fruits, while acetoin and aliphatic alcohols are volatiles commonly associated with fermented carbohydrate-rich substrates. Furthermore, fungal partners in all tripartite nursery pollination systems appear to consist of opportunistic fungi that are also fruit pathogens. The fungus *Rhizopus artocarpi* (= *Rhizopus stolonifer*) identified in the male inflorescences of *Artocarpus heterophyllus* (Gardner et al., 2018) is the primary cause for *Rhizopus* rot, which is a common disease in developing *Artocarpus* fruits (McMillan, 1974, 1986). In *A. integer*, its fungal partner *Choanephora* (Sakai et al., 2000) is similarly known to cause wet rots in various fruit crops (Kwon & Jee, 2005; Park et al., 2015). It is unclear whether the participation of these fungi in mutualistic relationships merely indicates their ubiquity, or whether fungal pathogens of fruits are predisposed to such pollination mutualisms. The fruity aromas and utilisation of fruit pathogens as fungal partners suggest that such systems likely arose from adopting mycetophagous insects associated with rotting fruits as the pollinators. These flowers might have tapped into the pre-existing communication channel used by mycophagous insects, which rely on cues of ripe fruits to locate the fruit rotting fungi associated with it. Indeed, the pollinator genus *Paraphloeostiba* has previously been found to associate with fermenting fruit (Kuschel, 1990; Thayer et al., 2003). Characterisation of the fruit fungal biome and arthropod assemblage on mature *M. hainanensis* fruits, as well as in common substrates infested with fungus in the vicinity could shed light on this hypothesis.

Given the similarities among tripartite nursery systems, we propose possible scenarios that may have given rise to tripartite nursery systems. (1) Tripartite nursery pollination might have evolved from an ancestral condition in which the flowers or early developing ovaries were regularly infected by fungi, causing a reduction in the reproductive fitness of the plant. Consequently, the plant adapted by shifting to mycetophagous pollinators that utilised flower-or ovary-infesting fungi as brood sites to keep the fungal infection at bay. Pollinator larvae probably impose the highest predation pressure on the fungi during the early stage of ovule development, since oviposition mostly takes place during anthesis. Indeed, the mycelia of fungal partners of *M. hainanensis* were seen infecting developing ovaries. In addition, the *Rhizopus* fungal partner of *Artocarpus heterophyllus* is also a major pathogen for its fruits and early developing ovaries (McMillan, 1974). This hypothesis might offer an explanation why tripartite systems reported so far all utilise fungal partners that are capable of infecting fruits and early developing ovaries. While this is compelling for the case of *M. hainanensis*, there is no evidence for pollinator oviposition on the female inflorescences of *Artocarpus* spp. (Sakai et al., 2000; Gardner et al., 2018), casting doubts on its validity. This postulation also highlights the importance of the larvae against the negative influence of fungal infection on ovaries. Control larvicidal experiments may be used to assess whether the removal of larvae significantly reduces fruit and seed sets of *M. hainanensis*. This hypothesis also predicts that the fungus-flower interaction predates the shift of pollinators in evolutionary timescale. (2) A second scenario is that the pollinator was a vector for fungal transmission. The shift of pollinator composition to these mycetophagous beetles would inoculate the flower with fungi. This scenario predicts that the shift of pollinators and fungus-flower interaction occurred at the same time in evolutionary timescale. (3) A third scenario is that the flower and pollinator have established a pre-existing bipartite nursery pollination mutualism, in which antagonistic fungal pathogens may occasionally infect the flowers. The pollinator progeny subsequently adapted by shifting their food substrate from the flowers to the fungi. In the bipartite nursery system between the flower of *Silene latifolia* and the pollinating seed-feeding moth *Hadena bicruris*, an antagonistic smut-disease fungus *Microbotryum violaceum* infects the flowers and developing fruits (Biere & Honders, 2006). It was shown that the pollinating moths were six times less likely to oviposit on fungus-infected flowers, suggesting that selection probably favoured the utilisation of uninfected flowers over shifting to feeding fungi. In addition, this scenario also predicted that the fungivorous behaviour of the pollinator larvae is a novel adaptation evolved after nursery pollination mutualism. However, in the case of *Meiogyne hainanensis*, the evolution of mycetophagous diet of the pollinator larvae presumably predated that of nursery pollination. The related lineages of the pollinators (monotomid beetles for *Mimemodes*; Omaliini beetles for *Paraphloeostiba*) are mostly reported to associate with mushrooms or fungus-rich substrates (Shavrin, 2010; Lee et al., 2020). Evidence based on available literature and our study thus does not seem to substantiate this postulation. Overall, the aforementioned hypotheses predicted different chronologies of evolutionary events, and would leave different evolutionary patterns on dated phylogenies. Additional evidence from molecular dating and ancestral character mapping for the plants, insects, and fungi may shed light on the evolutionary origin of tripartite nursery pollination systems.

Records for tripartite nursery pollination systems are rare in the literature, but likely underrepresented. In her review on brood-site pollination systems, Sakai (2002) summarised that most pollination systems that offer decomposing flowers as brood site rewards are found in the tropics, where efforts for pollination studies are less intense. Furthermore, the olfactory signalling of documented systems is mediated by branched-chain aliphatic esters (Gardner et al., 2018), which have been reported in the flowers of numerous lineages (Knudsen et al., 1993; Goodrich & Jürgens, 2018). Therefore, the utilisation of mycetophagous pollinators might be more common than the literature suggests. The inner petals of the congeneric *Meiogyne anomalocarpa* are also regularly infested with filamentous fungi while attached to the receptacle (Piya Chalermglin, pers. comm.), and might represent another study avenue. In the recent few decades, more studies have revealed that floral microbes, including fungi, have a significant role to play in pollination (Vannette et al., 2013; Schaeffer et al., 2017; Yang et al., 2019; Deng et al., 2024). There remains much to investigate on the influence of floral fungi on the reproductive success of flowers.

## Acknowledgements

We would like to express our greatest gratitude to Xiaojiang Hong, the vice president of the Administration of Hainan Tropical Rainforest National Park, the Bawangling National Forest Park, the Huishan Forest Nature Reserve Administration, and the Jianfengling National Forest Ecological Field Station in Hainan, for their permission to conduct field work. We would also like to thank Xing Chen and Qing Chen for their field assistance, and Laura Wong and Jessie Lai for their technical support. This research is funded by the Hong Kong Research Grants Council (HKU17112616), awarded to R.M.K.S.

